# Bacterial longevity requires protein synthesis and a stringent response

**DOI:** 10.1101/628826

**Authors:** Liang Yin, Hongyu Ma, Ernesto S. Nakayasu, Samuel H. Payne, David R. Morris, Caroline S. Harwood

**Affiliations:** Department of Microbiology, University of Washington, Seattle, USA; Department of Environmental Science and Engineering, Xi’an Jiaotong University, Xi’an, P.R. China; Biological Sciences Division, Pacific Northwest National Laboratory, Richland, WA, USA; Department of Biology, Brigham Young University, Provo, UT, USA; Department of Biochemistry, University of Washington, Seattle, USA

**Keywords:** *Rhodopseudomonas palustris*, ribosome, growth-arrest, translation, longevity, stringent response

## Abstract

Gram-negative bacteria in infections, biofilms and industrial settings often stop growing due to nutrient depletion, immune responses or environmental stresses. Bacteria in this state tend to be tolerant to antibiotics and are often referred to as dormant. *Rhodopseudomonas palustris*, a phototrophic α-proteobacterium, can remain fully viable for more than four months when growth is arrested. Here, we show that protein synthesis, specific proteins involved in translation and a stringent response are required for this remarkable longevity. Because it can generate ATP from light during growth arrest, *R. palustris* is an extreme example of a bacterial species that will stay alive for long periods of time as a relatively homogeneous population of cells and it is thus an excellent model organism for studies of bacterial longevity. There is evidence that other Gram-negative species also continue to synthesize proteins during growth arrest and that a stringent response is required for their longevity as well. Our observations challenge the notion that growth-arrested cells are necessarily dormant and metabolically inactive, and suggest that such bacteria may have a level of metabolic activity that is higher than many would have assumed. Our results also expand our mechanistic understanding of a crucial but understudied phase of the bacterial life cycle.

**IMPORTANCE:** We are surrounded by bacteria; but they do not completely dominate our planet despite the ability of many to grow extremely rapidly in the laboratory. This has been interpreted to mean that bacteria in nature are often in a dormant state. We investigated life in growth arrest of *Rhodopseudomonas palustris*, a proteobacterium that stays alive for months when it is not growing. We found that cells were metabolically active; they continued to synthesize proteins and mounted a stringent response, both of which were required for their longevity. Our results suggest that long-lived bacteria are not necessarily inactive but have an active metabolism that is well adjusted to life without growth.

## INTRODUCTION

Bacteria in nature often exist in viable but non-growing states without forming differentiated structures like spores (*1, 2*), but this crucial phase of the bacterial life cycle is underexplored. Factors that limit cell replication include nutrient limitation, competition within microbial communities and stresses like oxygen-depletion and extreme temperatures (*3–5*). Several bacterial pathogens, including *Vibrio cholera*, *Pseudomonas aeruginosa* and *Burkholderia pseudomallei* can survive for months and even years in a growth-arrested state in distilled water, sterilized seawater or basal salts medium in a laboratory setting (*6–9*). In keeping with this, drinking water is a known reservoir for bacterial pathogens (*10*). Non-growing pathogenic bacteria also occur in human infections and in this form present challenges for treatment because antibiotics tend to target processes such as cell-wall biosynthesis or DNA replication that are active in growing, but not non-growing cells. One example of a bacterium that is recalcitrant to antibiotic treatment *in vivo* is *Mycobacterium tuberculosis*, which forms latent infections that persist over many years in the absence of overt signs of growth (*11*). Then there are bacterial antibiotic persisters, a small subpopulation of non-growing or slowly-growing cells that develop in some bacterial infections and are tolerant to antibiotic treatment (*12–14*).

Studying non-growing bacteria is also important in environmental, industrial and other non-medical contexts. For example, non-growing bacteria are excellent biocatalysts because they can convert substrates that might be used for growth to value-added products. In past work we have studied production of hydrogen gas, a biofuel, by the phototrophic α *Rhodopseudomonas palustris.* When illuminated to allow ATP synthesis by photophosphorylation, non-growing cells diverted cell resources to hydrogen gas production and generated 300% more hydrogen gas than growing cells (*15*). We subsequently determined changes in metabolic fluxes and biomass composition that accounted for diversion of electrons to hydrogen gas in growth-arrested cells (*15*). We also found that non-growing *R. palustris* cells embedded in thin strips of latex film remained viable and produced hydrogen gas for over four months (*16*).

These results prompted us to develop *R. palustris* as a model for studies of bacterial longevity, and to ask the question: are there universal features of long-lived bacteria that we can uncover from studies of this proteobacterium? Several attributes make *R. palustris* a good model for addressing this question. First, it generates ATP and proton motive force from light, even when it is not growing (Figure 1). Also, *R. palustris* carries out photophosphorylation under anaerobic conditions and does not generate oxygen, thus avoiding complications of oxidative stress during growth arrest. A Tn-seq study of growth-arrested *R. palustris* identified 116 genes, which we call longevity genes, that are essential for long-term viability of non-growing cells, but are not required for growth (*17*). Among these are genes that are conserved in bacteria and associated with basic cellular processes including protein synthesis. Here, we carried out a physiological characterization of *R. palustris* in its growth-arrested, but viable state. We found that active translation and optimized ribosomes were critical for the longevity of *R. palustris,* as was guanosine polyphosphate [(p)ppGpp]. This goes against the conventional description of non-growth as state of metabolic inactivity often referred to as dormancy. We suggest that protein synthesis and other cellular activities are likely important for many bacteria to maintain viability when they are not growing.

**Figure 1.**
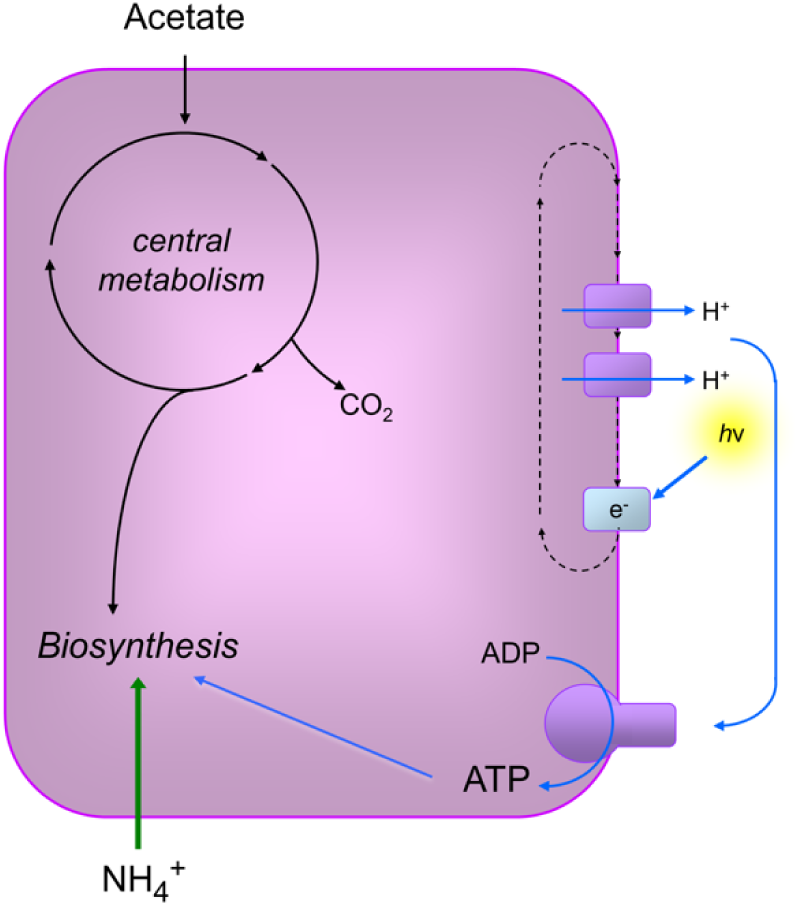
Metabolism of *R. palustris* during anaerobic growth in light. Acetate was used as the carbon source and ammonium as the nitrogen source for growth in this study. *R. palustris* generates ATP and proton motive force from light energy by cyclic photophosphorylation. Energy generation and carbon use are independent from each other. In this study growth arrest occurred when *R. palustris* had used all available acetate. In this circumstance cells can continue to generate ATP and proton motive force from light.

## RESULTS

### Conditions of growth arrest

In this study *R. palustris* was grown anaerobically in minimal salts medium in sealed glass tubes in light until growth arrest occurred due to carbon source depletion (*17*). Cultures were incubated post-growth arrest in the same tubes in which they were grown. Nitrogen was supplied in excess as ammonium and the carbon supplied was 20 mM sodium acetate. We took as day 0 of growth arrest, the time at which cells stopped growing as measured by a leveling off of the increase in the optical density (OD) of cultures. Cells grew to twice the OD when we supplied them with 40 mM acetate, confirming that growth arrest occurred due to carbon source limitation.

### Characterization of cell size

Upon growth arrest due to starvation for nutrients many bacteria decrease their size and change from a rod-shaped to coccid form (*6, 18–20*). Degradation of ribosomes that accompanies growth arrest typically results in a loss of total cellular protein and RNA, which likely contributes to cell shrinkage. To put our work on *R. palustris* into context with the large body of work on starvation-survival of heterotrophic bacteria, we measured cell size before and after growth arrest. As shown in Figure 2, cells in log phase and at 6 days post-growth arrest do not differ appreciably in length or overall morphology.

**Figure 2.**
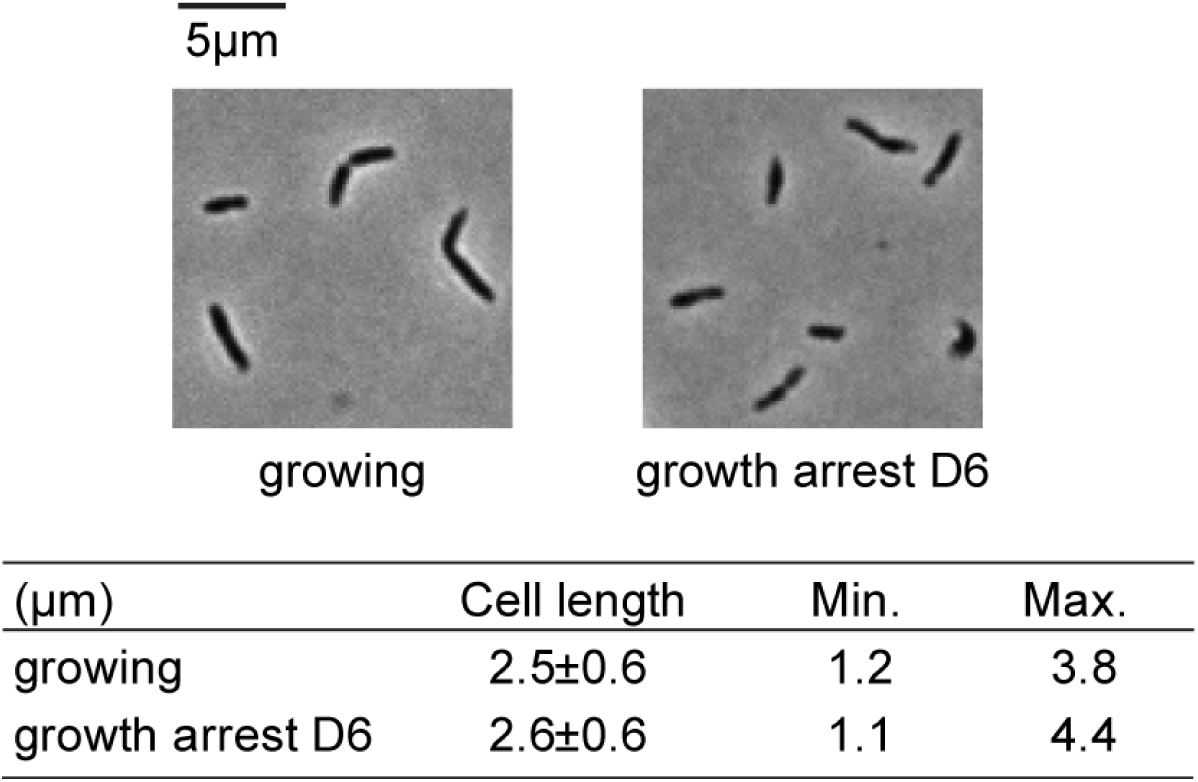
Comparison of lengths of *R. palustris* cells from growing cultures and from cultures at day 6 post growth arrest. Cells were photographed at 100X magnification with a Nikon Eclipse 80i digital microscope (n > 50).

### Growth-arrested *R. palustris* cells do not appear to undergo cycles of death and regrowth

To put the work reported here in context with another aspect of the starvation literature, we note that depending on the conditions of growth arrest some bacteria, the best studied of which is *E. coli*, undergo cycles of death and growth after entry into stationary phase (*1, 21*). Typically a fraction of cells survive long-term stationary phase by feeding on nutrients released from dead cells. This leads to heterogeneity among cells, making it difficult to carry out meaningful studies of growth-arrested bacteria at the population level. Also genetic heterogeneity develops in populations of growth-arrested bacteria as there is strong selection pressure for mutations that enable cannibalism (*22, 23*). In previous work we reported that populations of growth-arrested *R. palustris* cells were nearly 100% viable (*17*). To firmly establish that cells were truly growth arrested, rather than undergoing cycles of growth and death, we introduced an unstable plasmid expressing a gentamycin (Gm) resistance gene into wild-type *R. palustris*. When Gm is absent, the percentage of growing cells that retain this plasmid would be expected to decrease (*24*). On the other hand, a non-growing culture would be expected to retain the plasmid in Gm-free medium. When this strain (WT::pBBR^Gm^) was inoculated into Gm-free medium, about 60% of the cells had lost the plasmid by the time they reached stationary phase. When these cells were transferred into fresh Gm-free medium and successively transferred several times over a period of 20 days after reaching stationary phase, the plasmid continued to be lost. In contrast, growth-arrested cells retained the plasmid over the same period of time (Figure 3a). This suggests that most or all of the cells were indeed in a non-growing state. However, we cannot exclude that some small percentage of the cells underwent cycles of growth and death.

**Figure 3.**
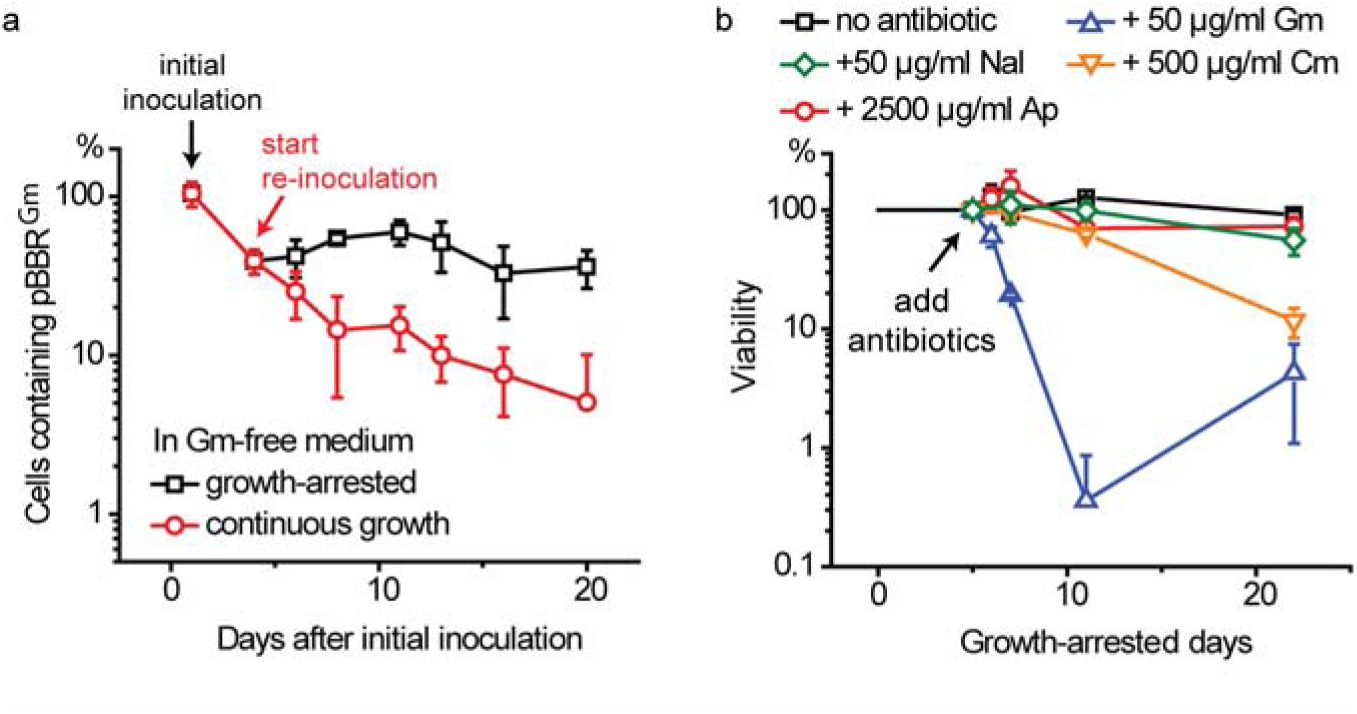
Growth-arrested *R. palustris* cells do not appear to undergo cycles of death and regrowth. a: Percentage of *R. palustris* cells that continue to maintain plasmid pBBR^Gm^ during continuous growth (re-inoculated in medium with fresh carbon source repeatedly) or during growth arrest (no treatment after the initial carbon source was depleted). Gm-free medium was used for this experiment. b: Growth-arrested *R. palustris* is sensitive to antibiotics targeting ribosomes. Antibiotics are added five days after growth arrest to independent cultures. Gm: gentamycin. Cm: chloramphenicol. Nal: nalidixic acid. Ap: ampicillin. Error bars indicate the standard deviation (n>3).

In further support of the conclusion that growth-arrested *R. palustris* does not undergo major cycles of growth and death, we found that non-growing cells were tolerant to the cell wall synthesis inhibitor ampicillin and the DNA synthesis inhibitor nalidixic acid at concentrations that prevented growth (Figure 3b and SI Figure 1). However, we also found that the viabilities of non-growing cells were compromised by protein synthesis inhibitors (Figure 3b and SI Figure 1).

### Long-lived, growth-arrested cells continue to synthesize proteins

Growth-arrested *R. palustris* was sensitive to Gm, an aminoglycoside antibiotic that affects protein synthesis by causing mistranslation, which results in accumulation of damaged proteins. We found that over 99% of growth-arrested cells lost viability over a few days when treated with Gm at a final concentration of 50 µg/ml (Figure 3b). Although cells died relatively rapidly, they were not as sensitive to Gm as growing cells, which died within a few hours after exposure to this antibiotic. We noticed that the viability of growth-arrested cells exposed to Gm at 20 d post-growth arrest tend to be higher than the number of viable cells at day 10 post-growth arrest (Figure 3b). It is possible that some cells developed resistance to Gm or that the Gm supplied may have become degraded, allowing for a small amount of growth on debris released from dying cells.

Chloramphenicol (Cm) belongs to another class of antibiotic that targets ribosomes. Although typically bacteriostatic, Cm was effective in killing growth-arrested *R. palustris* over a prolonged period of time (Figure 3b and SI Figure 1). These results suggest that active translation is important for *R. palustris* longevity.

Proteomics experiments revealed that after 6 days and 20 days of growth arrest, 289 and 525 proteins, respectively, were present at greater than two-fold higher levels than in growing cells (Figure 4a and SI Table 1). Among these were enzymes involved in the degradation of compounds, including benzoate and fatty acids (SI Tables 1 and 2), that are preferred carbon sources for *R. palustris*, suggesting that cells respond to growth arrest by increasing the translation of proteins that may aid in scavenging carbon. In the period between 6 and 20 days post growth arrest, 51 proteins of unknown function increased in abundance. Some proteins (163 and 310 proteins at days 6 and 20 post growth arrest) were diminished in abundance by two-fold or more following growth-arrest (Figure 4a and SI Table 1). This set included 53 out of 55 ribosomal proteins that were identified in our proteomics dataset. On average, ribosome proteins were at approximately 50% and 25% of their original levels after 6 days and 20 days of growth arrest (Figure 4b and 4c). We calculated the relative mass of these proteins per cell and from this calculated that ribosomes comprised 25.7% of the total protein mass in growing cells, which was reduced to 13.1% (49% reduction) and 6.8% (74% reduction) after 6 and 20 days of growth arrest (Figure 4c, SI Table 3). Not all translation related proteins were decreased in their abundances. For example, translation elongation factors made up ∼5% of the total protein from growing cells as well as growth-arrested cells (Figures 4d and 4e, SI Table 3). Similar patterns were also observed for proteins involved in aminoacyl-tRNA biosynthesis (tRNA charging) and for two previously identified longevity proteins YbeY and Era (*17*) that are involved in ribosome maturation (Figure 4d).

**Figure 4.**
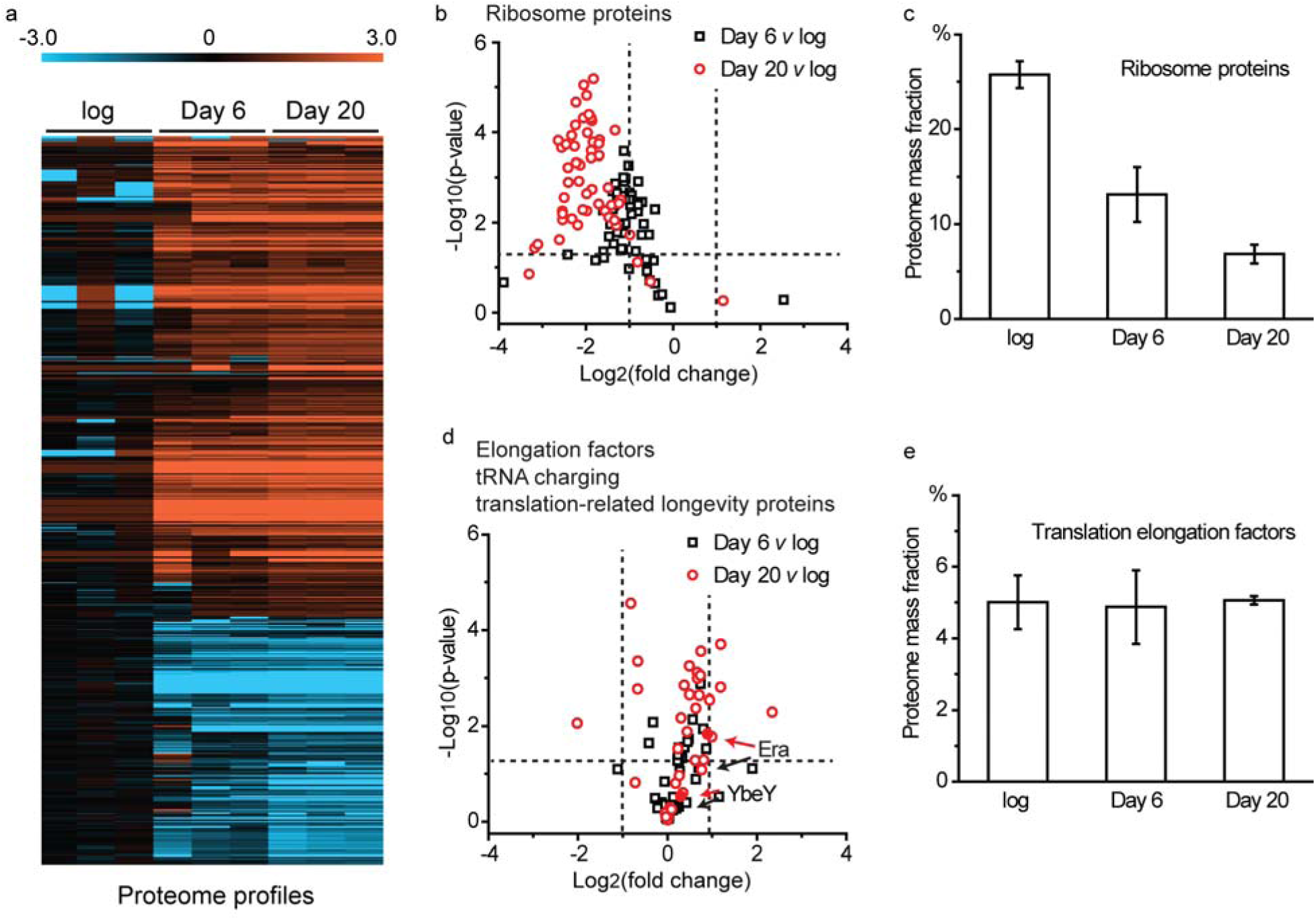
*R. palustris* protein abundances. Growing cells and growth-arrested cells are compared. a: Log_2_-fold changes in abundances of differentially abundant (p ≤ 0.05 by Student’s *t*-test) *R. palustris* proteins in growing cells (log), 6 days after growth arrest (Day 6) and 20 days after growth arrest (Day 20). The proteome profiles of three samples (n = 3) for each condition are shown (SI Table 1). b: The average relative quantities of 53 identified ribosome proteins at day 6 (black, Day 6) and day 20 (red, Day 20) growth arrest relative to the growing cells (log). The horizontal dashed line indicates the position of p = 0.05 (Student’s *t*-test), and the vertical dashed lines indicate the position of two-fold abundance changes. c: Ribosome proteins compared in growing cells (log) and at day 6 and day 20 post growth arrest as % mass faction of the proteome (SI Table 3). Error bars represent standard deviation. d: The average relative quantities of selected proteins at day 6 (black, Day 6) and day 20 (red, Day 20) growth arrest relative to the growing cells (log). The horizontal dashed line indicates the position of p = 0.05 (Student’s *t*-test), and the vertical dashed lines indicate the position of two-fold abundance changes. This panel includes proteins involved in tRNA charging (KEGG map00970, SI Table 2), translation elongation factors (SI Table 2) and two previously identified longevity proteins (YbeY and Era). e: Translation elongation factors (SI Table 3) compared in growing cells (log) and at day 6 and day 20 post growth arrest as % mass fraction of the proteome. Error bars represent standard deviation.

Finally, to directly examine protein synthesis in growth-arrested *R. palustris*, we constructed a LacZ-reporter driven by a P_hirI_ promoter that can be induced with phenylacetyl-homoserine lactone (PA-HSL) (SI Figure 2). After 6 days of growth arrest, PA-HSL was added to the culture and samples were collected after 2 additional days of incubation. The negative control without added signal showed a base level of LacZ expression. Compared to this, LacZ acitvity was ∼70% higher when the PA-HSL signal was added to induce lacZ expression (SI Figure 2). This observation directly demonstrates that *R. palustris* has the ability to synthesize proteins during growth arrest.

### Characterization of protein synthesis apparatus in long-lived cells

We isolated *R. palustris* ribosomes by sucrose gradient centrifugation and found that the ribosome profile of growing *R. palustris* cells included 30S, 50S and 70S populations, as is typical for growing bacteria (Figure 5a). There was also a small peak corresponding to a 100S population, which is likely a dormant form of ribosomes, as has been seen in other bacteria (*25, 26*). The ribosome profile of growth-arrested *R. palustris* cells resembled that of growing cells (Figure 5a and 5b), but the quantity of the 70S population was lower at day 6 and even lower at day 25 post growth arrest, suggesting that cells degraded some fraction of their ribosomes after they stopped growing. This is consistent with our proteomics data (Figure 4b) and fits the notion that the total rate of protein synthesis is expected to be much lower in growth-arrested cells compared to growing cells. Ribosome degradation is often accompanied by the non-specific degradation of rRNA (*27–30*). However, even after 120 d of growth arrest, cells retained intact species of 23S and 16S rRNA, indicating that cells continuously repaired and replaced ribosomes albeit at probably a low rate after they stopped growing (Figure 5c, SI Figure 3). The relative levels of aminoacylated (or charged) and free forms of tRNAs are important for promoting and inhibiting translation (*31*). We used Northern blot analysis to estimate the percentage of tryptophan-charged tRNA_trp_ (Trp-tRNA_trp_) relative to tRNA_trp_. In growing cells, ∼60% of tRNA_trp_ was in the charged Trp-tRNA_trp_ form (Figure 5d and 5e). This level is comparable to that reported in other bacteria (*32*). The percentage of Trp-tRNA_trp_ increased to ∼85% upon growth arrest and was maintained at this level for ∼20 days (Figure 5d and 5e).

**Figure 5.**
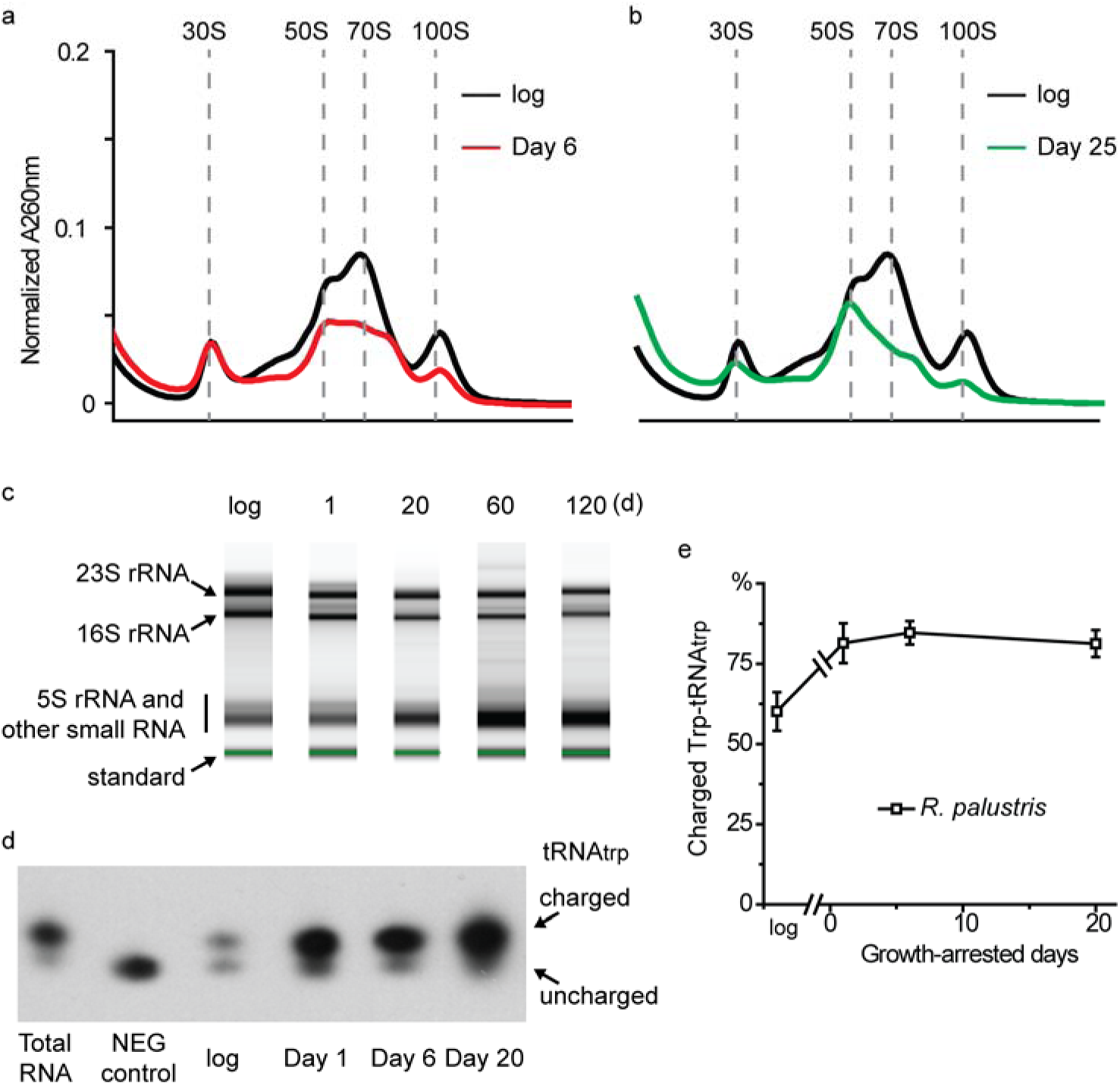
Translation is intact and active in growth-arrested *R. palustris*. a: The ribosome profile of cells at 6 days (Day 6) post growth arrest is shown and compared to ribosomes from growing *R. palustris* (log). b: The ribosome profile of cells at 25 days (Day 25) post growth arrest is shown and compared to ribosomes from growing *R. palustris* (log). c: Total RNA was purified and analyzed from growing and growth-arrested *R. palustris* cells. The same amount of RNA was loaded in each lane. The quantification of replicated experiments is shown in SI Figure 3. The standard is the internal standard of the RNA ScreenTape system used (Agilent). d: The charged and uncharged species of tRNA_trp_ are shown here with a representative Northern blot. The aminoacylated (charged) and deacylated (uncharged) species of tRNA_trp_ were separated with an acid urea gel and probed with a tRNA_trp_ probe. e: Quantification of the charged and uncharged species of tRNA_trp_ as analyzed in Figure 5d. Error bars represent the standard deviation (n>3).

### Two proteins identified as essential for *R. palustris* longevity are critical for ribosome integrity in growth-arrested cells

Among the longevity genes that we have identified in *R. palustris* (*17*), *ybeY* and *era* are predicted to encode proteins that interact with each other and the S11 ribosome protein during ribosome maturation (*33–37*). *ybeY* or *era* deletion mutants have compromised longevity but grow normally (*17*). Here we found that the ribosome profiles of growing cells were similar between the wild type and these mutants (Figure 6a). All had distinctive 30S, 50S and 70S populations, although the 50S subunit was slightly more abundant in the mutants. The ribosome profiles of growth-arrested *ΔybeY* and *Δera* mutant cells were noticeably different from that of the wild type, however. A prominent 50S subunit peak was seen but the amounts of 30S subunits and 70S ribosomes were greatly diminished (Figure 6a).

**Figure 6.**
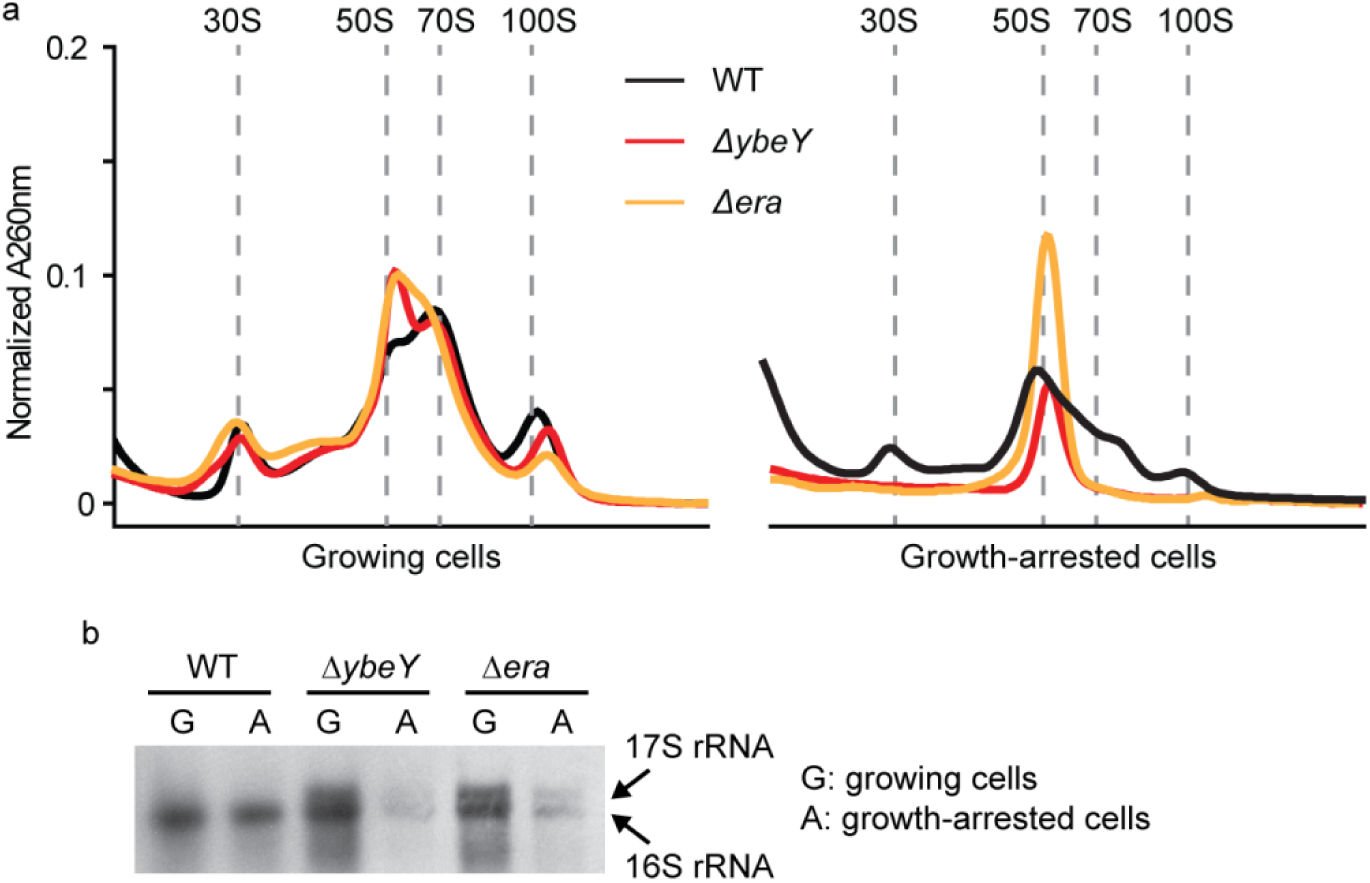
YbeY and Era are important for ribosome maintenance during growth arrest. a: Deletion of either *ybeY* or *era* had little effect on ribosomes from growing *R. palustris*, while the ribosomes from growth-arrested cultures of *ybeY* and *era* deletion mutants lacked 30S subunits. b: *R. palustris* total RNA was analyzed by Northern blot with a probe targeting 16S rRNA. In growing cells, both Δ*ybeY* and Δ*era* mutants accumulated 17S rRNA but maintained substantial levels of 16S rRNA. In growth-arrested cells, both 17S rRNA and 16S rRNA species were depleted.

As predicted based on studies of their *E. coli* homologs (*33–36*), *ΔybeY* and *Δera* strains were defective in 16S rRNA maturation. An extra band immediately above the 16S rRNA was present in gels of RNA from both mutants (SI Figure 3). Northern blot experiments confirmed that the extra band was 17S rRNA, a precursor of 16S rRNA (Figure 6b). In growth-arrested cells, both 16S rRNA and 17S rRNA species were severely depleted in the mutated strains (Figure 6b and SI Figure 3). This is likely connected to the defective 30S ribosome subunit production shown in Figure 6a, and the compromised longevity of *R. palustris*. We do not know why the 17S and 16S rRNAs were so much lower in abundance in growth-arrested vs. growing cells (Figure 6b and SI Figure 3). It is possible that factors associated with these mutations coupled with the stress of growth arrest lead to their degradation. *ybeY* or *era* deletion mutants had wild-type levels of charged tRNA-Trp (SI Figure 4), suggesting that tRNA aminoacylation is not directly coordinated with the process of ribosome maturation.

As shown in Figure 4b, there was no significant change in the abundance of YbeY and Era proteins in growth-arrested cells. Since the core ribosomal proteins were decreased 2 to 4-fold in abundance (Figure 4b), the relative abundance of YbeY and Era was increased on a per ribosome basis following growth arrest. To test whether we could change the effects of YbeY on longevity by modulating its relative abundance, we constructed and expressed a YbeY variant predicted based on work in *E. coli*, to interact weakly with ribosomes. In *E. coli*, the interaction between YbeY and ribosomes is partially disrupted by replacement of a conserved aspartate with an arginine (*37*). When we made the corresponding YbeY_D118R_ change in *R. palustris*, the strain was compromised in its longevity to a similar degree as the *ΔybeY* mutant (Figure 7a). When we overexpressed YbeY_D118R_ *in trans* (*ΔybeY*::pBBP-*ybeY*_D118R_) however, the longevity defect was complemented (Figure 7a). The quantity of 16S rRNA was similar to that of the wild-type, and no 17S rRNA was observed during growth arrest (Figure 7b). The ribosome profile of the growth-arrested *ΔybeY*::pBBP-*ybeY*_D118R_ -complemented strain differed from that of growth-arrested wild-type cells, especially in the quantities of 50S and 70S ribosomes (Figure 7c). However, growth-arrested cells could apparently tolerate these differences when challenged in longevity assays (Figure 7a). These results suggest that it may be helpful for growth-arrested cells to have relatively more YbeY protein compared to growing cells and that this promotes 30S ribosome subunit assembly in this condition.

**Figure 7.**
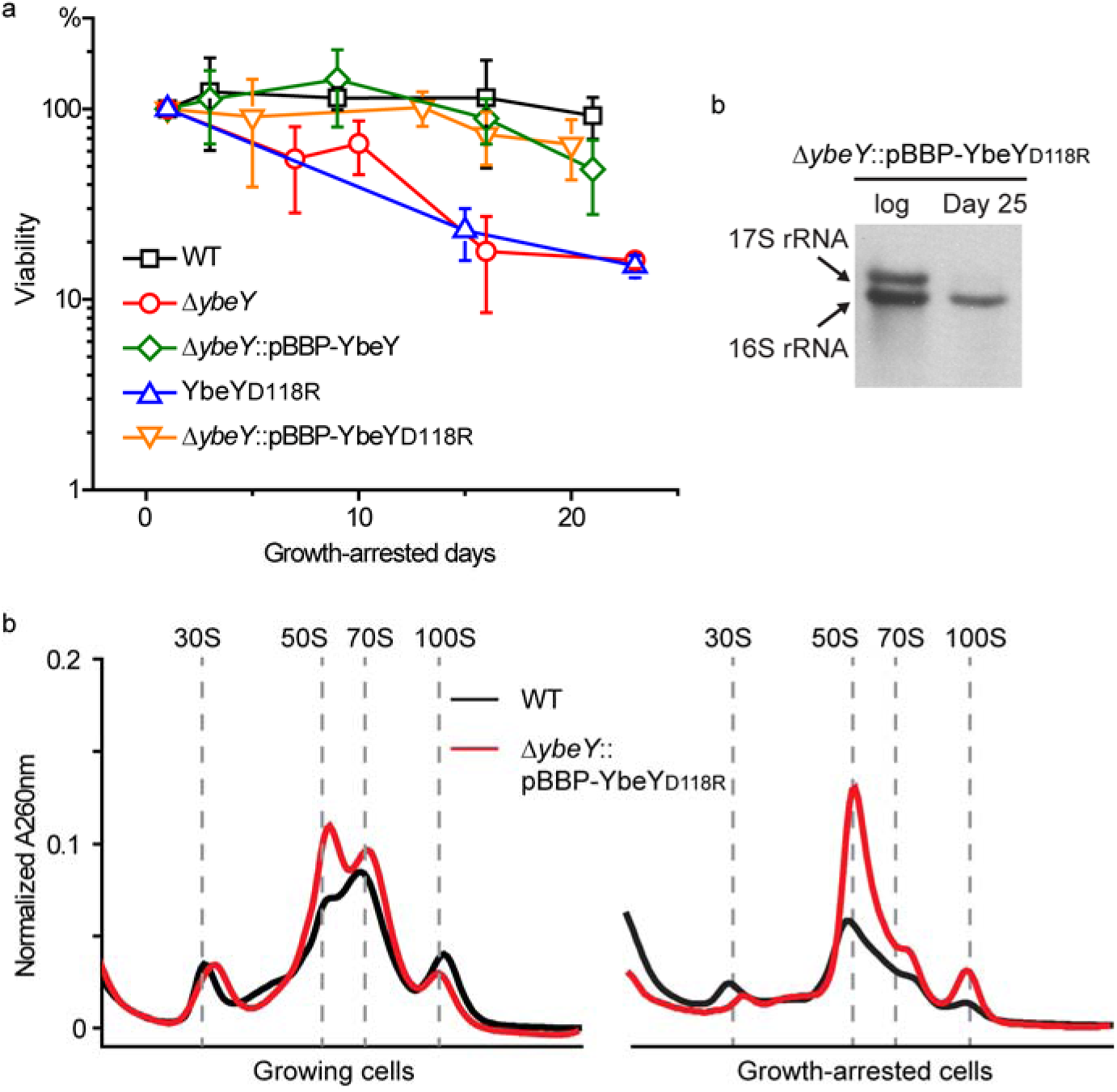
Overexpression of YbeY_D118R_ complemented the longevity defect of a Δ*ybeY* mutant. a: Longevity of different *R. palustris* strains. Deletion of the *ybeY* gene compromised the longevity of growth-arrested cells (Δ*ybeY*), and this was complemented by expressing this gene *in trans* (Δ*ybeY*::pBBP-YbeY) (*17*). A *ybeY*_D118R_ mutant had a longevity defect similar to that of the Δ*ybeY* deletion mutant. Overexpression of *ybeY*_D118R_ *in trans*, restored the longevity of a Δ*ybeY* mutant (Δ*ybeY*::pBBP-YbeY_D118R_). Error bars indicate the standard deviation (n>3). b: *R. palustris* total RNA was analyzed by Northern blot with a probe targeting 16S rRNA. In the growth-arrested cells of Δ*ybeY*::pBBP-*ybeY*_D118R_, only 16S rRNA appeared and no 17S rRNA was detected. c: The ribosomes from Δ*ybeY*::pBBP-YbeY_D118R_ were analyzed on a sucrose gradient. In the growth-arrested cells, 30S, 50S and 70S ribosomes were clearly seen, though the abundance of the 50S subunit is higher than that of WT.

### The stringent response is important for longevity

Our results suggest that translation is reduced but still maintained in growth-arrested cells. A well-known mechanism that contributes to this is the stringent response mediated by guanosine polyphosphate [(p)ppGpp] (*38, 39*). In many bacteria, the level of (p)ppGpp increases upon growth arrest and this inhibits the transcription of ribosomal genes, although the mechanism by which this is achieved varies among bacteria (*38–40*). As with other bacteria, we detected little to no (p)ppGpp in exponentially growing *R. palustris*. However, the level of (p)ppGpp increased sharply as growing cells reached stationary phase and remained at relatively high levels after growth arrest (Figure 8a).

**Figure 8.**
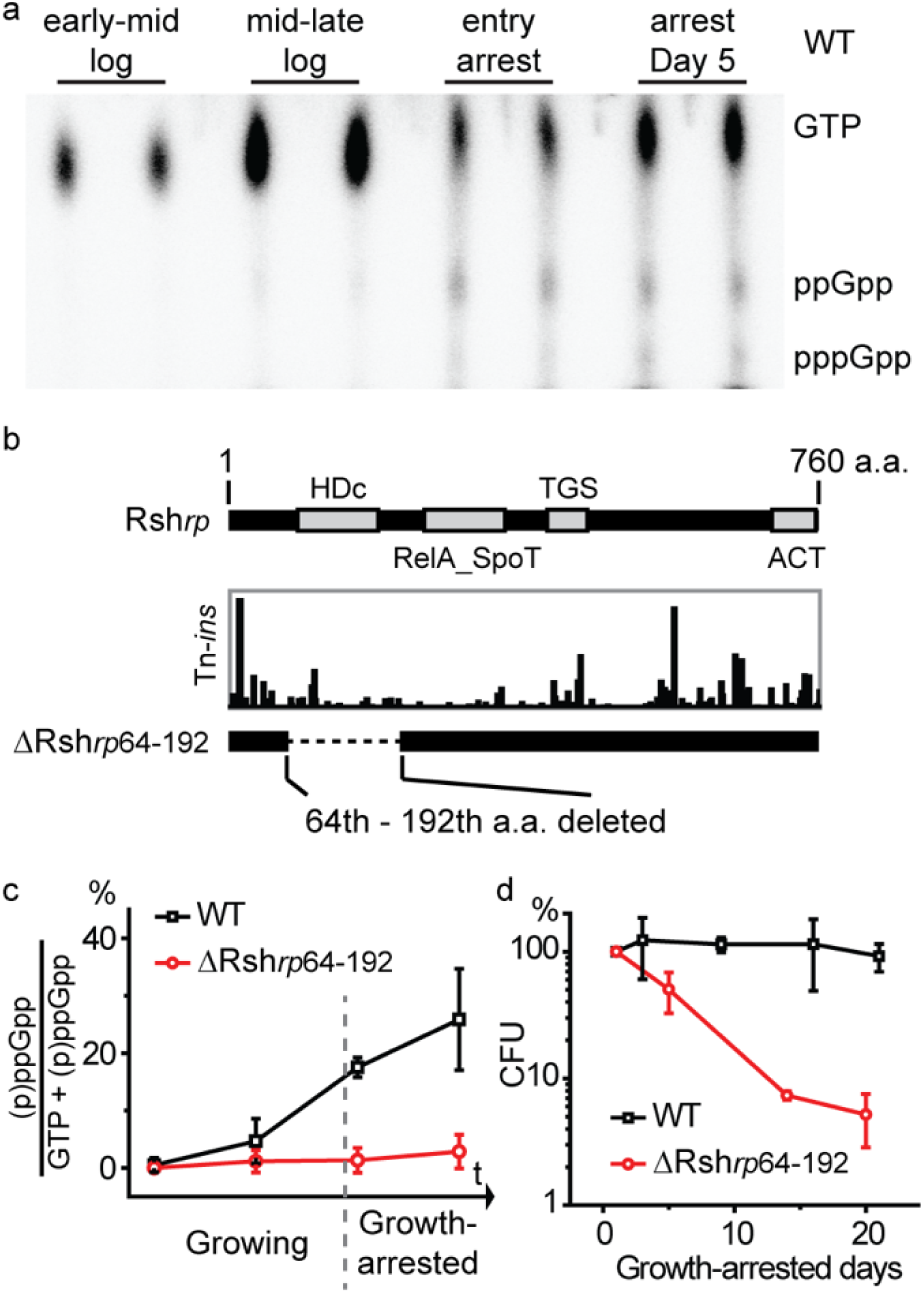
Growth-arrested cells with very low levels of (p)ppGpp have compromised longevity. a: The level of (p)ppGpp was low in exponentially growing *R. palustris* and increased during growth-arrest. b: The domain structure of Rsh*_rp_* (top). Locations and relative quantities of Tn insertions in Rsh*_rp_* (*17*). A ΔRsh*_rp_*_64-192_ mutant was made by deleting the sequence encoding the 64th-192th amino acids of Rsh*_rp_* from *R. palustris.* c: (p)ppGpp synthesis was severely compromised in the ΔRsh*_rp_*_64-192_ strain. The time points shown correspond to the time points shown in Figure 8a, representing early-mid log phase, mid-late log phase, entry into growth-arrest and day 5 following growth-arrest. d: The longevity of Δrsh*rp*64-192 was severely compromised. Less than 10% of cells remained viable after 2 weeks of growth-arrest. Error bars indicate the standard deviation (n>3).

Unlike *E. coli*, but like most other bacteria outside the γ- and β-proteobacteria, *R. palustris* has a single bifunctional (p)ppGpp synthesis/hydrolysis *rsh* gene, here called *rsh_rp_*, that has multiple domains including a hydrolase (HDc) and RelA_SpoT domain (Figure 8b). Our Tn-seq results indicated that this gene (*rpa2693*) was essential for *R. palustris* longevity, but not for growth (*17, 41, 42*). However, we were unable to delete the full-length *rsh_rp_* gene. A subsequent inspection of the locations of the Tn-insertions in this gene in our bank of Tn mutagenized cells revealed that there were relatively few insertions in its RelA_SpoT domain (Figure 8b). When we made domain specific deletions in Rsh*_rp_*, we obtained a hydrolase domain deletion mutant (ΔRsh*_rp_*_64-192_, Figure 8b) that produced very little (p)ppGpp during growth or growth arrest. (Figure 8c). We do not know why this deletion affected (p)ppGpp synthesis in this way, but one idea is that it destabilized or altered the structure of the protein so that its ability to synthesize (p)ppGpp was compromised. The ΔRsh_*rp*64-192_ mutant grew with a generation time of 9.4 ± 0.6 h, compared to a wild type 6.4 ± 0.2 h generation time, a growth rate difference of about 30%. In contrast to this relatively modest effect on the growth rate, the ΔRsh*_rp_*_60-189_ mutation had a severe effect on longevity. As shown in Figure 8d, the mutant survived poorly in growth arrest.

## DISCUSSION

We found that illuminated *R. palustris* responds to growth arrest by reducing ribosome abundance as is typical of most bacteria when they slow their growth rate. Growth-arrested cells continued to synthesize proteins, some at elevated levels relative to growing cells, over the period of 20 days that we followed protein composition by proteomics (SI Table 1) and it is likely that they repaired and replaced some portion of their original ribosome inventory over this time period. The levels of most proteins involved in tRNA charging and translation elongation and were at similar levels in growing and growth-arrested cells (Figure 4d and SI Table 2), which is consistent with the notion from a previous report that elongation rates are maintained as bacteria slow down their growth-rates even towards zero growth (*43*). In addition, a tRNA that we followed was almost fully charged 20 d into growth arrest (Figure 5d and 5e). Not only does protein synthesis continue after growth arrest, but it is essential for viability of non-growing cells as indicated by their sensitivity to Gm and Cm (Figure 3b, SI Figure 1).

There are reports of continued protein synthesis in other species of bacteria in growth arrest, the more convincing of which come from studies in which production of fluorescent proteins in single cells was used as a proxy for protein synthesis. Antibiotic persisters in early stages of formation, as well as non-growing *E. coli*, *Mycobacterium* and *Salmonella* have all been inferred to carry out protein synthesis (*44–47*). These results suggest that maintenance of some level of protein synthesis may be required for viability of many non-growing bacteria. From this it follows that it would make sense to prioritize development of antibiotics that target protein synthesis in growth-arrested cells. This will likely require understanding in detail what happens to bacterial ribosomes following growth arrest. Bacterial ribosomes are known to be differentially post-translationally modified in stationary phase cells (*48*). Two longevity genes, *ybeY* and *era*, were essential for normal ribosomes to form in non-growing *R. palustris* (Figure 6a), and their encoded proteins increased abundance on a per ribosome basis during growth arrest (Figure 4b). Era is a well-studied bacterial GTPase (*36*), and genes for it and YbeY, are present and conserved in most bacteria (*34*). Our data are consistent with studies of the corresponding *E coli* proteins showing that YbeY and Era are involved in rRNA processing in *R. palustris*. When we looked at rRNAs formed by Δ*ybeY* and Δ*era* mutants that were actively growing, it was apparent that growing cells and growth-arrested cells had the same defect in 17S rRNA processing (Figure 6b), but only the growth-arrested cells had a physiological phenotype. This highlights a feature of at least some *R. palustris* longevity proteins, which is that they are functional in growing cells as well as in non-growing cells, but it is only when the additional demands of growth arrest on cellular functionality are imposed that essentiality becomes apparent in growth arrest. Our work on Era and YbeY has led us to conclude that optimized ribosomes are a feature of long-lived cells.

In early stationary phase and when in growth arrest, *R. palustris* synthesizes increasing amounts of (p)ppGpp, indicating that cells mount a stringent response. The gene *rpa2693*, which we call *rsh_rp_*, is among the genes identified by Tn-seq to be essential for longevity (*17*). A Δ mutant did not produce detectable (p)ppGpp and the viability of this mutant decreased rapidly after growth arrest (Figure 8c and 8d). There is good evidence that (p)ppGpp is also important for development of antibiotic persisters (*49*) and for long term survival of latent and growth-arrested *M. tuberculosis* (*50, 51*). A great deal still needs to be learned about how (p)ppGpp signaling is activated in growth-arrested *R. palustris*. Studies of (p)ppGpp in different species of bacteria have generally not followed the levels of this nucleotide deep into stationary phase. *E. coli* has been shown to have elevated levels of (p)ppGpp only in early stationary phase, with (p)ppGpp levels returning to baseline later in stationary phase. This is very different from the situation in *R. palustris* where high levels of (p)ppGpp are maintained for at least five days post-growth arrest. Uncharged tRNAs are understood to be the classic signal for activation of RelA and up-regulation of (p)ppGpp synthesis in *E. coli* and other gamma proteobacteria (*40*). But in *R. palustris*, tRNA_Trp_ charging is even higher in growth arrest than during growth (Figure 5d and 5e). It could be that one or several tRNA species other than tRNA_Trp_ become uncharged during carbon starvation, which could potentially trigger (p)ppGpp synthesis in a fashion similar to the case of *E. coli*. However, it is also known for that the exact triggers of (p)ppGpp synthesis can vary between species (*40, 52, 53*). For example, amino acid starvation that typically leads to uncharged tRNA species is insufficient to trigger (p)ppGpp accumulation in *Rhizobium meliloti* (*54, 55*), *Caulobacter crescentus* (*56*) and *Rhodobacter sphaeroides* (*57*). A nitrogen-related phosphotransferase system (PTS^NTR^) has been found to be important for stimulating (p)ppGpp production in *C. crescentus* in response to nitrogen limitation (*58, 59*), but the mechanism by which carbon limitation, the condition studied here, may stimulate (p)ppGpp accumulation α-proteobacteria remains unclear (*56*).

Other future areas of investigation are to understand why (p)ppGpp is essential for *R. palustris* longevity and to identify its target genes and proteins. Based on work with other bacteria it is very likely that (p)ppGpp interacts with RNA polymerase to inhibit transcription of genes encoding ribosomal RNAs and proteins (*38–40*). But this would need to be done in a very controlled fashion because some new ribosome biogenesis is very likely to be required for *R. palustris* longevity in growth arrest. (p)ppGpp has recently been shown to bind to Era from *Staphylococcus aureus* and *E. coli* to inhibit its GTPase activity (*60–62*), and a similar effect on Era from *R. palustris* might influence the biogenesis and integrity of its ribosomes.

Virtually all bacteria probably stop growing at one time or other due to nutrient limitation in their native niches and it is also likely that that they employ diverse strategies to stay alive. However, there also may be evolutionarily conserved core survival strategies that all bacteria use. We suggest that optimized ribosomes, a stringent response and protein synthesis are part of this core strategy The primary reason for the extreme longevity of *R. palustris* is likely its ability to continuously generate ATP by photophosphorylation when it is in growth arrest, because growth-arrested cells incubated in dark do not survive nearly as well (*17*). All bacteria need ATP to stay alive. There is plenty of evidence in the literature suggesting that ATP, obtained by scavenging resources from the environment or from degradation of internal energy stores or cell biomass is necessary to support the viability of diverse non-spore forming growth-arrested bacteria (*20*). A recent study also suggested that varying ATP levels per cell might underlie the heterogeneity in populations and lead to non-growing persisters (*63*). Protein synthesis and (p)ppGpp synthesis are core features that underlie the extraordinary longevity of *R. palustris*. Both of these processes are highly energy-consuming and crucial for all bacteria. To understand how *R. palustris* integrates these processes and utilizes its ATP during growth-arrest could be a key to understand how non-growing bacteria in general stay alive.

## METHODS

### Bacterial strains, growth and incubation conditions

*R. palustris* strain CGA009 was used as the wild-type for this study. Anaerobic cultures were grown under illumination at 30 °C using a defined media PM with 20 mM sodium acetate or 10 mM sodium succinate as the carbon source (*64*). The longevity of *R. palustris* cultures was determined by counting colony-forming units as described previously (*17*). In-frame deletions of *R. palustris* genes were created by using suicide vector pJQ200SK as described before (*65*). For *in trans* expression in *R. palustris*, the target genes were cloned into plasmid pBBPgdh and the plasmids were mobilized into *R. palustris* by conjugation using *E. coli* strain S17-1 (*66*). Media were supplemented with 100 µg/ml of gentamicin for *in trans* complementation experiments.

### Construction of inducible LacZ-reporter and LacZ assays

To construct the inducible LacZ-reporter, we assembled a *lacZ* fragment and a *hirR*-P_hirI_ fragment (*67*) into a pBBP plasmid backbone (*66*) (SI Figure 2). The promoter P_hirI_ responds specifically to PA-HSL signal (*67*). At day 6 of *R. palustris* growth arrest, PA-HSL was added into the culture at final concentration of 1 µM. After an additional ∼48 h of incubation, 5 ml of culture was collected and resuspended in 2 ml of Z-buffer (60 mM Na_2_HPO_4_, 40 mM NaH_2_PO_4_, 12.5 mM NaCl, 1 mM MgSO4 and 40mM 2-mercaptoethanol, pH 7.0). The samples were sonicated at a frequency of 20∼40 kHz for 3 cycles (20 s sonication and 20 s cooling on ice). Cell lysis was cleared by centrifugation at 4 °C, 15,000 rpm for 15 min, and the supernatant was collected for subsequent β-galactosidase assays. To start the reaction, 200 µl supernatant was mixed with 40 µl of 4 mg/ml ONPG at 30 °C, and Abs_420nm_ was monitored using a Biotek H1 plate-reader. The level of LacZ was calculated as the V_max_ of the reaction normalized by total protein concentration of the supernatant.

### Ribosome purification

Ribosomes from *R. palustris* cells were isolated following a method adapted from a previous study (*68*). About 35 ml of culture was grown to the desired state and poured over a 10-ml ice sheet (fast-cool). The ice sheet melted within 10 min and the cells were collected by centrifugation at 4000 rpm for 15 min. The cell pellet was resuspended in 2 ml of Buffer A (20mM HEPES-KOH, pH 7.5, 30mM NaCl, 8mM MgCl_2_) with 1 µl/ml RNasin (Promega). Three hundred microliter of 0.1 mm Zirconia/Silica beads (Biospec Products) were added to every ml of sample. The samples were lysed with a Mini-Beadbeater-24 (Biospec Products) at 3500 rpm for 1 min at 4 °C, kept on ice for 1 min and the cycle was repeated 4 more times. The micro-beads and cell debris were removed by centrifugation at 15000 rpm for 10 min. The lysates were further clarified by spinning at 30,000 g in Beckman TL-1000 ultracentrifuge for 30 min, followed by pelleting ribosomes at 100,000 g for 4 hours. The pellet was resuspended with Buffer A overnight on ice.

### Sucrose gradient analysis

About five A_260_ units of ribosomes (less than 200 µl in volume) were applied on top of 11 ml of a sucrose gradient (7-47%). The samples were centrifuged through the gradient at 39,000 rpm for 4.5 h with a SW41 rotor, and then were separated with a Brandel gradient pump. The sucrose gradient was made with Buffer A.

### Ribosomal RNA analysis

Total RNA was purified from 3-5 ml of *R. palustris* culture using an miRNeasy Mini Kit. The rRNAs were then analyzed with an RNA ScreenTape system (Agilent). To probe 16S rRNA by Northern blot, 1 µg of total RNA was heated at 70 °C for 10 min and immediately chilled on ice. The samples were then separated on a 1% agarose gel with 1X TBE buffer. After electrophoresis, RNA was transferred to an Amersham Hybond-XL membrane (GE Lifesciences) with a downward nucleic acid transfer system overnight using 20X SSC buffer. A probe (5’-TCGTGCCTCAGCGTCAGTAATGGCCCAGTGA) labeled with DIG was used to detect 16S rRNA following the manufacturer’s instruction (DIG, Roche).

### Charging analysis of tRNA_Trp_

Total RNA was isolated with QIAzol lysis reagent according to the manufacturer’s instructions (Qiagen). The dried RNA pellet was dissolved and stored in 10 mM sodium acetate, pH 4.5. A deacylated control sample was prepared by incubating 10 µg of total RNA with 1 mM EDTA and 100 mM Tris-HCl pH 9.0 at 42 °C for an hour (*69*). Northern blot and acid-urea gel electrophoresis were performed essentially as described previously (*32, 69*). Loading dye (0.01% xylene cyanol, 0.01% bromophenol blue, 8 M Urea, 1 mM EDTA and 10 mM sodium acetate, pH 5.0) was mixed with sample at 1:1 ratio. Samples were loaded on a 12% acid-urea gel of 20 cm and then were run at 15 W for 12 h at 4 °C. Transfer of RNA to an Amersham Hybond-XL membrane (GE Lifesciences) was performed in 0.5* TBE buffer at 650 mA for 30 min 4 °C. The aminoacylated and the deacylated tRNA_Trp_ were detected using a DIG-labeled probe (5’-GGTTTTGGAGACCGGTGCTCTACCAATTGAG) following the manufacturer’s instruction (DIG, Roche).

### Determination of cellular (p)ppGpp levels

*In vivo* (p)ppGpp labeling and detection were performed as described with some modifications (*70*). Anaerobic *R. palustris* cultures were grown in 1.5 ml PM medium with 100 µCi/ml of Na_2_H^32^PO_4_ (PerkinElmer-NEX011001MC) added at the time of inoculation. At the desired time points, 200-300 µl of culture was collected and extracted with 8 M formic acid. Supernatants of the extracts were spotted on PEI-cellulose TLC plates (Sigma-Aldrich) and separated with 1.5 M KH_2_PO_4_ (pH 3.4). The plates were then dried and visualized by phosphor imaging. The level of (p)ppGpp was calculated by the ratio of (ppGpp + pppGpp) / (GTP + ppGpp + pppGpp).

### Proteomic analysis

Biological triplicates of cells were lysed, digested with trypsin and analyzed by liquid chromatography-tandem mass spectrometry as described in (*71*). Collected data were analyzed using MaxQuant software (v 1.5.5.1) (*41, 72*) for peptide/protein identification and quantification. Identification of peptides was done by searching against the *R. palustris* ATCC BAA-98 (strain CGA009) sequences from the Uniprot Knowledgebase (downloaded on August 30, 2016). The search was performed first with a larger parent mass tolerance (20 ppm) to identify peptides and re-calibrate peptide masses. After re-calibration, the parent mass tolerance was set to 4.5 ppm for a second round of search. During the searches oxidation of methionine as a variable modification, and trypsin digestion was required in both peptide termini. Identifications were filtered with 1% false-discovery rate at peptide-spectrum match and protein levels. The function “matching between runs” was checked to detect peptide peaks in LC-MS/MS runs that they were not fragmented. Label-free quantification (LFQ) values were used for quantitative analysis. This method considers only the most reliably detected peptides and normalizes according to the total signal of the run. It is, therefore, ideal for comparing individual proteins in different samples (*73*). For absolute quantification, intensity-based absolute quantification (iBAQ) (*74*) measurements of individual proteins were sequentially normalized by the protein molecular mass and total mass in the sample, resulting in relative protein mass compared to the entire cell proteome. Proteins were considered differentially abundant with a p-value ≤0.05 by T.test considering two tails and equal variance. The mass spectometry data have been deposited to the ProteomeXchange Consortium via the PRIDE partner repository with the dataset identifier PXD013729.

## Acknowledgements

The authors thank Amy Schaefer, Kathryn Fixen, Yasuhiro Oda and Yanning Zheng for helpful discussions, and Karl Weitz for technical assistance with the mass spectrometry analysis. Mass spectrometry data was generated in the Environmental Molecular Science Laboratory, a U.S. Department of Energy (DOE) national scientific user facility at Pacific Northwest National Laboratory (PNNL) in Richland, WA. Battelle operates PNNL for the DOE under contract DE-AC05-76RLO01830.

## Abbreviations

Tn: transposon

## Author contribution

L.Y. and C.S.H. designed the research. L.Y. and H.M. performed the antibiotics assays with *R. palustris*. L.Y. made the *R. palustris* mutants. L.Y. grew the *R. palustris* cells. S.H.P. and E.S.N. performed proteomic analysis. L.Y. and D.R.M. carried out the *in vitro* analysis of ribosome. L.Y. carried out the analysis of rRNA and tRNA. L.Y. performed the analysis of (p)ppGpp. All authors analyzed the data. L.Y., E.S.N. and C.S.H. wrote the paper. All authors contributed to the revision of the manuscript.

## SI Figure and Table Legends

**SI Figure 1.** The responses of *R. palustris* to various antibiotics at different growth stages. Antibiotics are added to growing cultures or five days after growth-arrest. Cm: chloramphenicol. Nal: nalidixic acid. Error bars indicate the standard deviation (n>3). For growing cells treated with Nal, the number of viable cells was determined by A_660_ and a representative result is shown (n > 3).

**SI Figure 2.** The *hirR* gene is expressed with a GDH (*rpa0944*) promoter and LacZ protein is expressed with P_hirI_ promoter. *R. palustris* culture was grown to Day 6 of growth-arrest and the PA-HSL signal was added into the culture. After two days, the samples were collected and the level of LacZ was analyzed as stated in the Method session. With the signal added, the LacZ level was ∼70% higher comparing to the negative control.

**SI Figure 3.** rRNA levels in different *R. palustris* strains at different growth-stages. The quantities of 16S, 17S and 23S rRNA are determined using an RNA ScreenTape system (Agilent) and normalized against the levels of 23S rRNA in growing cells of wild-type *R. palustris*. An unpaired *t* test was performed with at least three replicates (n.s.: not significant; *: p < 0.05; **: p < 0.01; ***: p < 0.005; ****: p < 0.001, as determined by Student’s *t*-test).

**SI Figure 4.** Trp-tRNATrp remained highly charged in the growth-arrested cells of Δ*ybeY* and Δ*era* strains.

**SI Table 1.** Proteome of R. palustris in log phase and at 6 days and 20 days post growth arrest.

**SI Table 2.** Categories (KEGG) of proteins with changes in abundances in growth-arrested compared to growing *R. palustris*.

**SI Table 3.** Relative mass % of individual *R. palustris* proteins compared to the whole cell proteome in log phase cells and in cells at 6 days and 20 days post growth arrest.

