## Supplementary figures and images for "Bacterial longevity requires protein synthesis and a stringent response"

### SI Fig 1

#### Slide 1
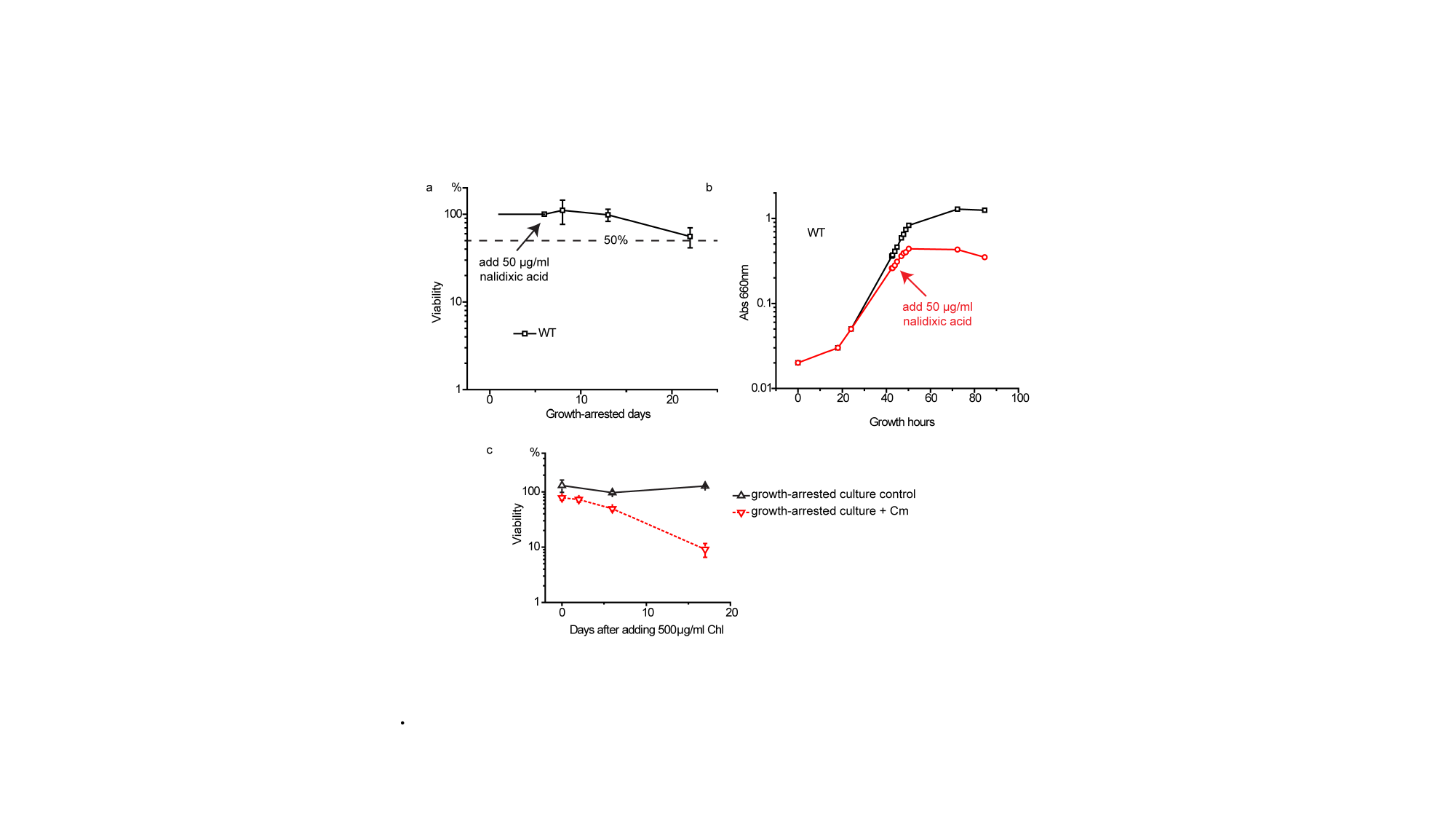

.

### SI Fig 2

#### Slide 1
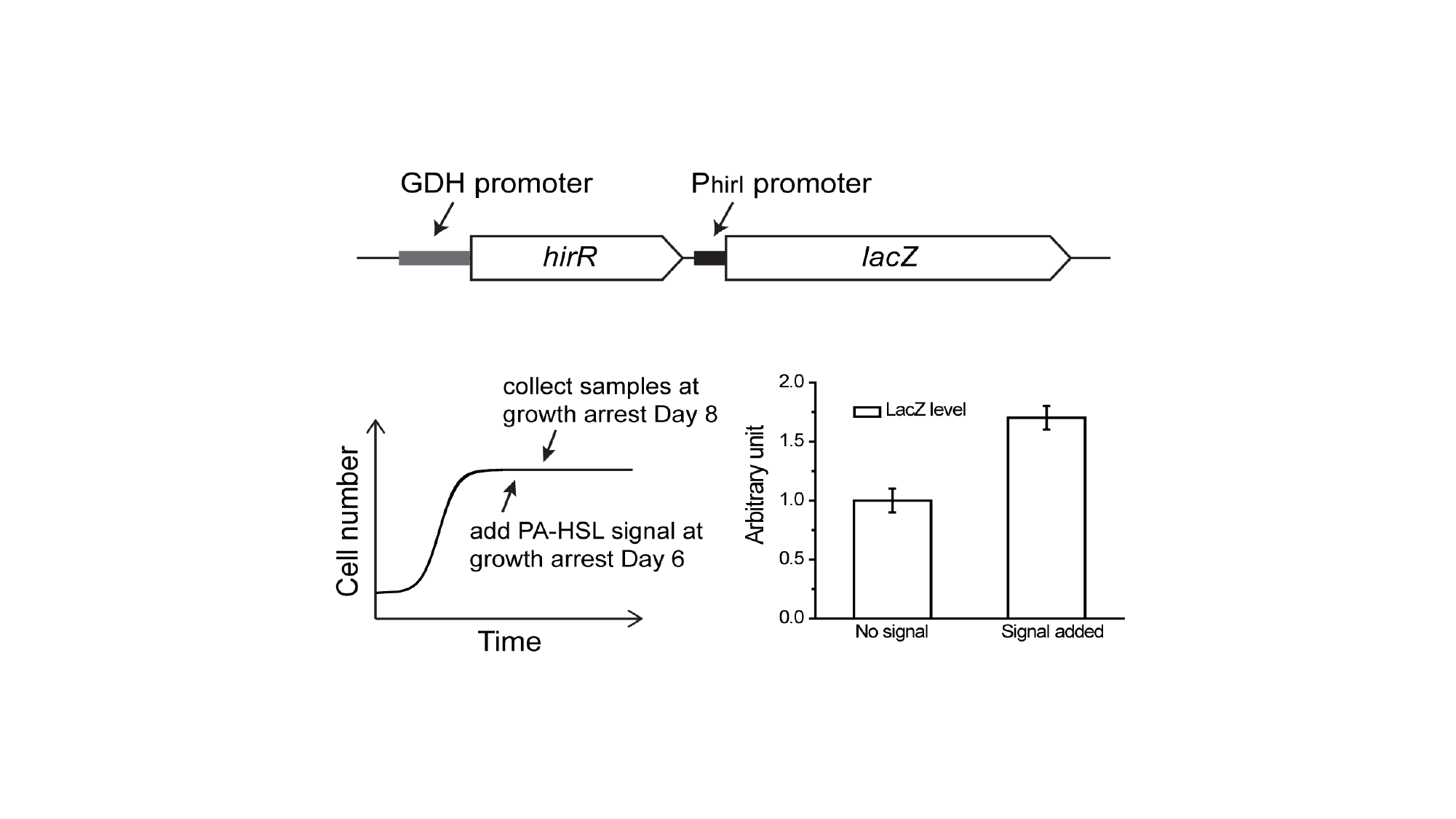

### SI Fig 3

#### Slide 1
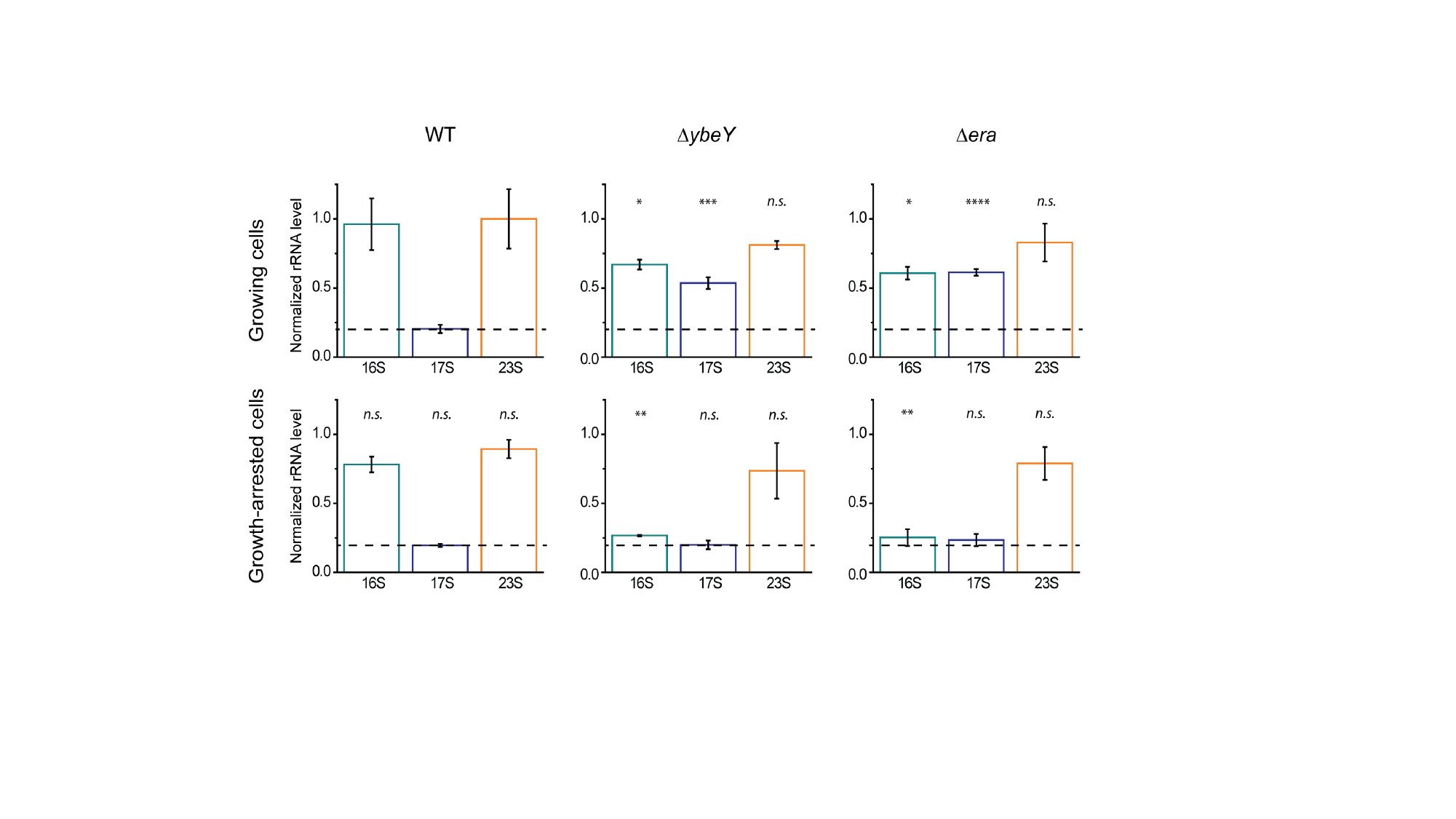

### SI Fig 4

#### Slide 1
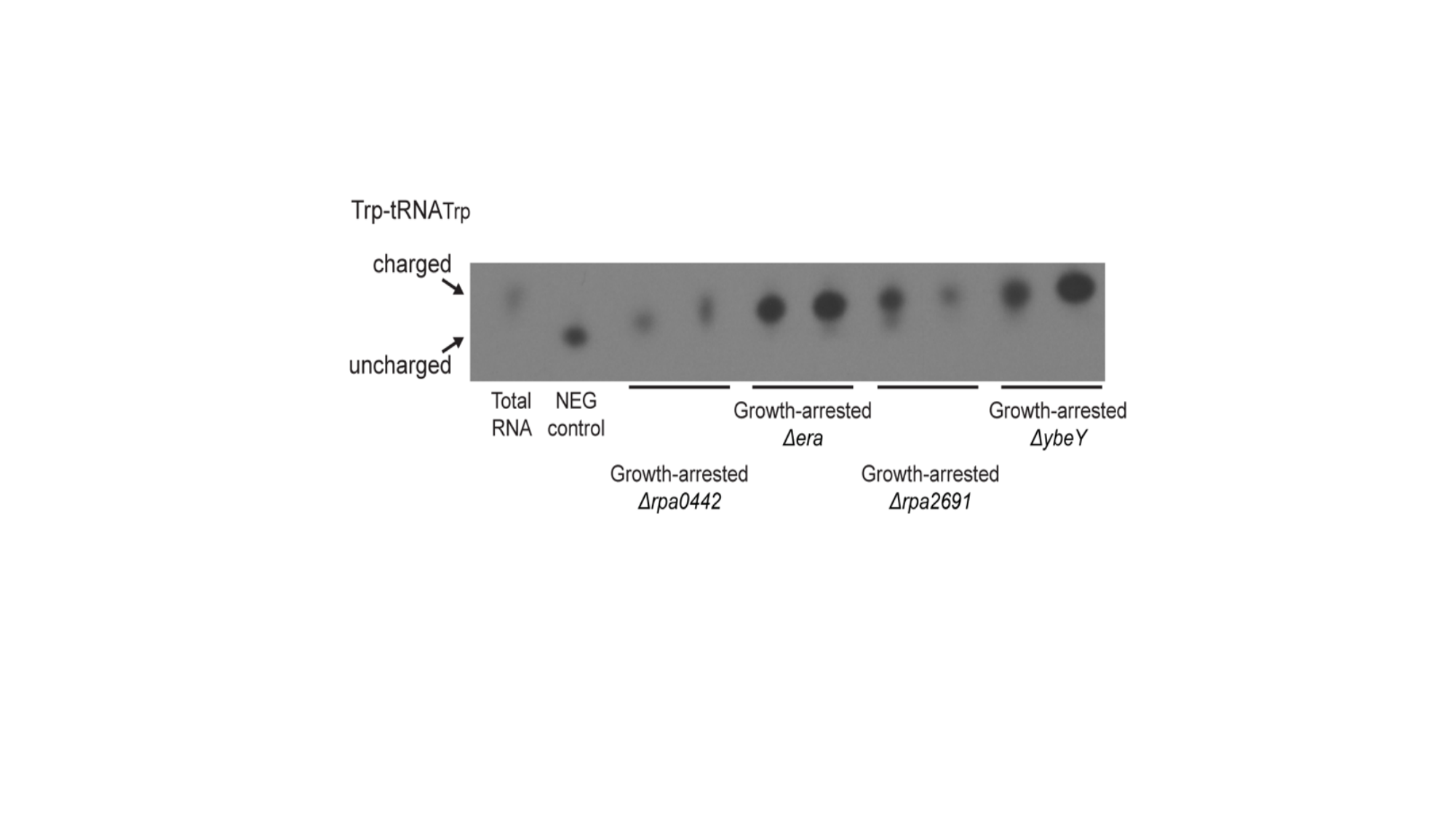
